# Defining the components of the miRNA156-SPL-miR172 aging pathway in pea and their expression relative to changes in leaf morphology

**DOI:** 10.1101/2021.08.22.457055

**Authors:** J.K. Vander Schoor, V. Hecht, G. Aubert, J. Burstin, J.L. Weller

## Abstract

The timing of developmental phase transitions is crucial for plant reproductive success, and two microRNAs (miRNA), *miR156* and *miR172*, are implicated in the control of these changes, together with their respective *SQUAMOSA promoter binding-like* (*SPL*) and *APETALA2* (*AP2*)-like targets. While their patterns of regulation have been studied in a growing range of species, to date they have not been examined in pea (*Pisum sativum*), an important legume crop and model species. We analysed the recently-released pea genome and defined nine *miR156*, 21 *SPL*, four *miR172*, and five *AP2-like* genes. Phylogenetic analysis of the *SPL* genes in pea, *Medicago* and *Arabidopsis* confirmed the eight previously defined clades, and identified a ninth potentially legume-specific *SPL* clade in pea and *Medicago*. Among the *PsSPL*, 14 contain a *miR156* binding site and all five *AP2-like* transcription factors in pea include a *miR172* binding site. Phylogenetic relationships, expression levels and temporal expression changes identified *PsSPL2a/3a/3c/6b/9a/9b/13b/21, PsmiR156d/j* and *PsmiR172a/d* as the most likely of these genes to participate in phase change in pea. Comparisons with leaf morphology suggests that vegetative phase change is unlikely to be definitively marked by a change in leaflet number. In addition, the timing of *FT* gene induction suggests that the shift from the juvenile to the adult vegetative phase may occur within fourteen days in plants grown under inductive conditions, and calls into question the contribution of *miR172/AP2* to the floral transition. This work provides the first insight into the nature of vegetative phase change in pea, and an important foundation for future functional studies.

## Introduction

There are two major phase changes or transitions in plant development after germination: vegetative phase change (juvenile vegetative to adult vegetative) and reproductive phase change (adult vegetative to adult reproductive). Collectively the timing of these transitions are crucial for reproductive success and if they occur too early or late the plant may lack the energy required to reproduce effectively through compromise of leaf function, or may reproduce when environmental conditions are not favourable.

The morphological variations that accompany vegetative phase change (VPC) across plant species are diverse. Changes in leaf morphology, particularly in the early juvenile phase, provide an illustration of this. For example, in maize (*Zea mays*) the first leaf is small and elliptical with a blunt end, but the subsequent leaves are lanceolate and pointed (Bongard-Pierce et al., 1996). In soybean (*Gylcine max*), the first two leaves produced after the cotyledons are simple and opposite, whereas the third and fourth leaves are trifoliate and frequently distichous (Yoshikawa et al., 2013). Some plants also exhibit several distinct intermediate forms in the juvenile and adult stages. In the strongly heteroblastic tree from New Zealand, *Pseudopanax crassifolius*, eight different types of leaves are produced across the seedling, juvenile and adult phases of growth (Gould, K.S., 1993).

Dramatically heteroblastic species such as ivy, certain eucalypts and acacias can create the misleading impression that phase change is discrete and simple. However, research into many species including the genetic “model” *Arabidopsis* have found the changes to be more subtle. It seems that variation in vegetative organs during shoot ontogeny is in general more likely to be continuous and gradual, with a trajectory of change in vegetative traits that starts in the juvenile phase and continues through to senescence. It follows that the transition from the juvenile to the adult vegetative phase must be occurring somewhere along that continuum before reproduction and that the nature and duration of this period of transition may be different across plant species. This makes comparisons and identification of common features difficult, but raises the question as to whether there is any universal underlying genetic and molecular mechanism that might generate these diverse morphological and physiological manifestations of VPC.

It is now considered that two microRNAs (miRNAs), *miR156* and *miR172*, and their targets, *SQUAMOSA promoter binding–like* (*SPL*) and *APETALA2-like* (*AP2-like*) transcription factors respectively, are central components in a conserved pathway for regulation of phase change (Fig. 1). These were first identified in *Arabidopsis* and maize (Chuck et al., 2007a; Wu and Poethig, 2006), and have subsequently been shown to play similar roles across a range of herbaceous and woody plants (Xie, 2006; Wang et al., 2011; Zhang et al., 2011; Shikata et al., 2012; Bergonzi et al., 2013; Zhou, et al., 2013; Silva et al., 2019; Zhou et al., 2021).

**Figure 1.**
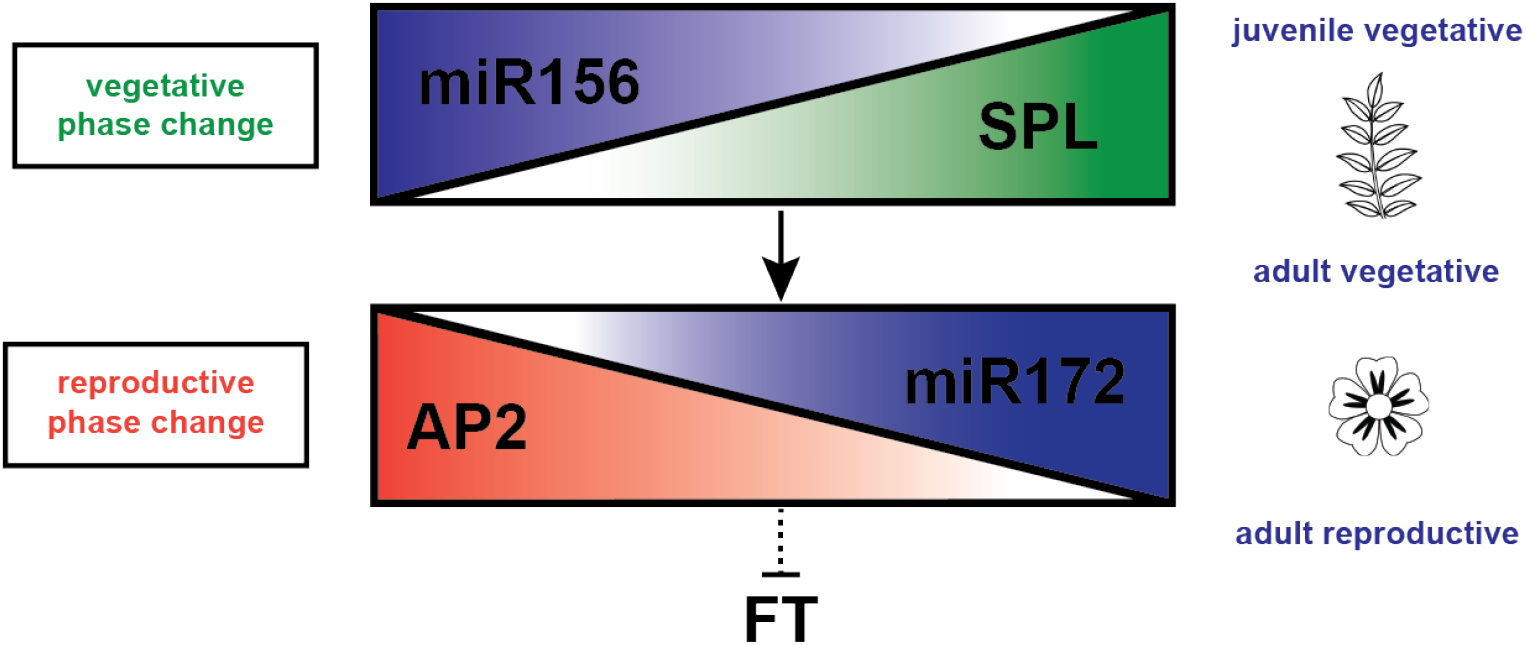
The *miR156-SPL-miR172-AP2* expression changes controlling phase change in plants. *MiR156* expression is high in the juvenile stage of plant development, directly suppressing expression of target *SPL* genes. As the plant ages, *miR156* levels decrease and *SPL* expression increases, which in turn upregulates *miR172*. At this stage the plant shifts into the adult vegetative phase and acquires competence to reproduce. *AP2-like* transcription factors act to suppress florigen (*FT*) genes and prevent flowering. As *miR172* expression increases, *AP2-like* genes are suppressed, and their repression of *FT* expression is in turn relieved, allowing the plant to initiate flowering and enter the adult reproductive phase.

The SPL proteins constitute a family of plant-specific transcription factors that regulate a wide range of biological functions within plant growth and development including leaf, flower and fruit development, coordinating the timing of development, and response to stress (Klein et al., 1996; Cardon et al., 1997; Stone et al., 2005; Manning et al., 2006; Wu and Poethig, 2006; Usami et al., 2009). *SPL* genes were first discovered in snapdragon (*Antirrhinum majus*) and identified on the basis of their regulatory interaction with the floral meristem identity gene *SQUAMOSA* (Klein et al., 1996). They have since been shown to be present in the model species *Arabidopsis* and many other diverse plant species from green algae and moss to cereals, vegetable and industrial crops and trees (Cardon et al., 1997; Arazi et al., 2005; Guo et al., 2008; Salinas et al., 2012; Li and Lu, 2014; B Wang et al., 2015; Zhang et al., 2015; Tripathi et al., 2017). They vary in size but are characterised by the highly conserved DNA binding domain (SQUAMOSA promoter binding protein, or SBP domain) always encoded in the first two exons (Guo et al., 2008). The SBP domain is approximately 75 amino acids in length and consists of a DNA binding domain containing two zinc binding (Zn1 and Zn2) sites and a nuclear localisation signal (NLS) that partly overlaps with the DNA-binding domain at the C terminus (Cardon et al., 1999; Yamasaki et al., 2004).

Many *SPL* genes contain a *miR156* binding site either within the last exon of the coding sequence or in the 3’UTR. For example 10 of the 16 *Arabidopsis SPL*, 11 of the 19 rice and 10 of the 15 tomato *SPL* contain *miR156* binding sites (Rhoades et al., 2002; Xie, 2006; Salinas et al., 2012). MiRNAs are 20-24 nucleotides in length and regulate gene expression at the post-transcriptional level by mRNA cleavage or by inhibiting translation (Bartel, 2004; Schwab et al., 2005). They originate from intergenic regions and in plants exhibit near-perfect sequence complementarity to their target mRNAs (Rhoades et al., 2002; Schwab et al., 2005; Nozawa et al., 2012). In *Arabidopsis*, the functional significance of the *miR156-SPL* interaction has been well documented. Overexpression mutants of *miR156*-resistant *SPL* result in accelerated vegetative phase change and premature flowering, similar to mutants in which *miR156* action is blocked through target mimicry, where a non-cleavable RNA forms a non-productive interaction with the complementary *miR156* target sites in *SPL* genes (Franco-Zorrilla et al., 2007). Conversely, *SPL* loss-of-function mutations delay VPC (Cardon et al., 1997; Wu and Poethig, 2006; Gandikota et al., 2007; Schwarz et al., 2008; Wang et al., 2009; Xu et al., 2016).

Specific *SPL* genes in turn promote the transcription of *miR172* (Wu et al., 2009), which target a number of *AP2-like* transcription factors that repress flowering in part by inhibition of *FT*. The interaction between *miR172* family and its *AP2* or *AP2-like* targets is deeply conserved in flowering plants and has been functionally tested in a number of species including rice, maize, barley and potato (Aukerman and Sakai, 2003; Lauter et al., 2005; Chuck et al., 2007b; Nair et al., 2010; Lee et al., 2014). When *miR172* is overexpressed in *Arabidopsis* (35S::miR172), plants undergo precocious vegetative phase change and early flowering. Similarly, plants in which the *miR172* targets *TOE1* and *TOE2* have been mutated show earlier expression of adult traits. Conversely, both the *miR172* knockout mutant and the *35S::TOE1* plants showed a delay in all phases of plant development (Aukerman and Sakai, 2003; Jung et al., 2007; Wu et al., 2009).

Legumes are a widely diverse group of plants from woody trees and climbing vines to annual herbaceous food crops, so it is not surprising that there is also enormous diversity in vegetative morphology. Woody acacia species can show distinct foliar dimorphism between juvenile and adult states (Hackett, 1985; Wang et al., 2011). Among the annual crop legumes, beans and soybean have two simple opposite leaves after germination, followed by alternate trifoliate leaves, whereas peas and lentils have markedly reduced scale leaves before producing compound leaves of increasing complexity in the adult plant. To date, the only study in crop legumes looking at developmental changes in shoot morphology associated with expression levels of *miR156-SPLs-miR172* is in soybean, where it is postulated that the first two simple leaves represent the juvenile phase, the third and fourth trifoliate leaves mark a transitional period and the adult vegetative stage is from the 5^th^ leaf onwards (Yoshikawa et al., 2013).

Despite the prominence of pea as a crop and historical model, there has been only a limited attempt to understand vegetative phase change in this species (Wiltshire et al., 1994), and no examination of the *miR156-SPLs-miR172-AP2* gene families or their potential relevance to developmental changes in leaf morphology. With the recent release of the pea transcriptome (Alves-Carvalho et al., 2015) and genome (Kreplak et al., 2019), a comprehensive analysis is now possible. This study presents the characterisation of these families with respect to structure, phylogenetic relationships, and developmental expression profiles, providing a solid foundation for future functional studies.

## Results

### The genomic location and sequence analysis of the pea *SPL* genes

To identify *SPL* genes in pea, the 16 *Arabidopsis SPL* were blasted against the pea reference transcriptome and genome (Alves-Carvalho et al., 2015; Kreplak et al., 2019) using BLASTp (Altschul et al., 1990). To confirm and compare the pea sequences with a second closely-related legume species, the *Arabidopsis* genes were also queried against the *Medicago truncatula* transcriptome/genome (Tang et al., 2014). Some *Medicago SPL* sequences have been reported previously (Preston & Hileman, 2013; Aung *et al*., 2015b; Gao *et al*., 2016; Wang *et al*., 2019), but since there is a lack of consensus on nomenclature in these studies, and the only account of the full family recently used a neighbour-joining algorithm for the phylogenetic analysis, we have here proposed a new unified nomenclature (based on maximum-likelihood analysis; see below) (Supplementary Table 1). 21 *SPL* sequences in pea were identified and located on the 7 pea chromosomes (Fig. 2A, Table 1). The *PsSPL* were evenly distributed across the genome with 3-4 genes per chromosome, except for the single *SPL6a* on Chr2. One gene, *SPL8*, was found to be located on an unmapped scaffold, but the genomic position was inferred from the surrounding genes and comparison with the syntenic region of chromosome 8 in *Medicago* (Supplementary Table 2).

**Figure 2.**
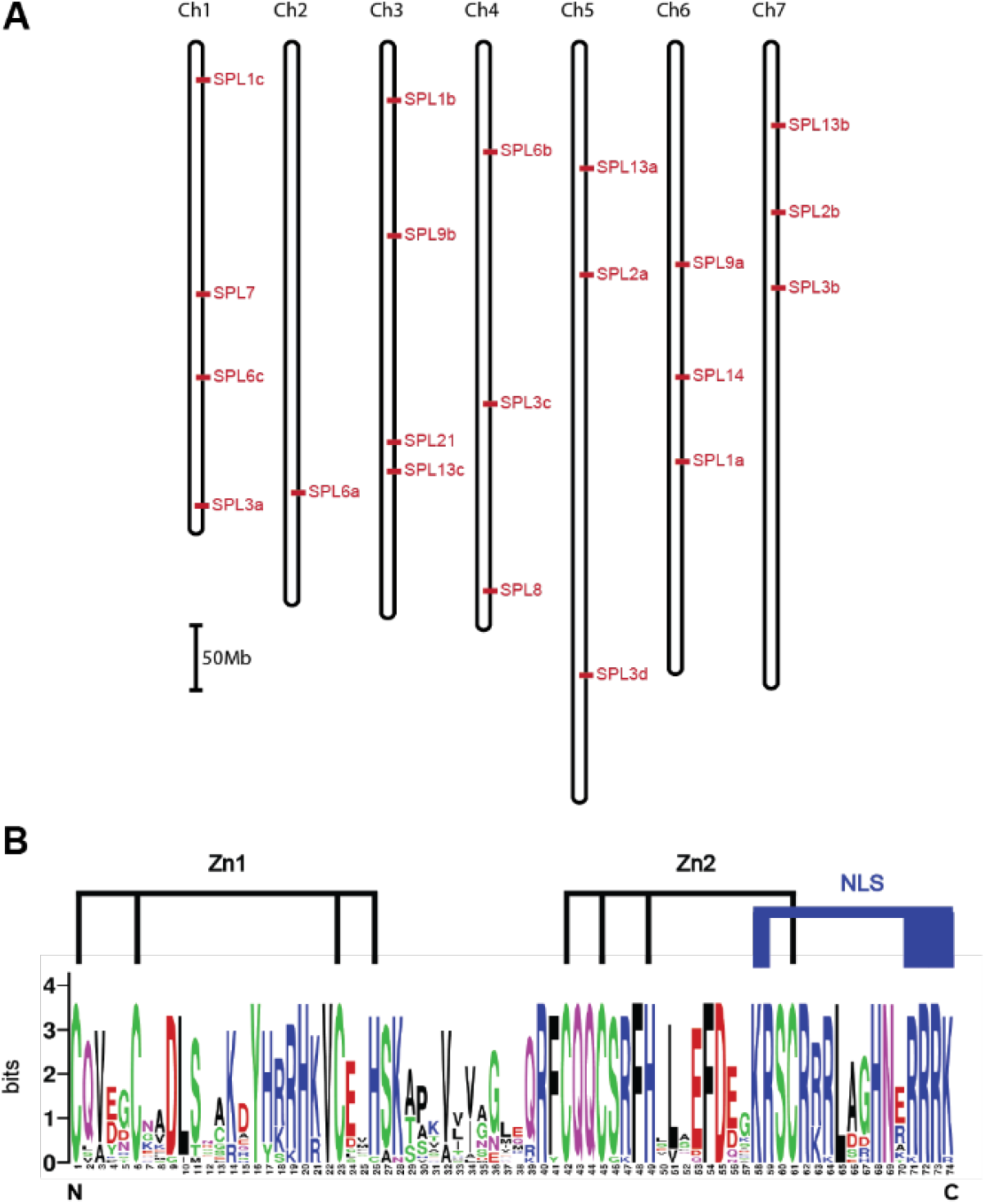
Location and sequence conservation of the *SPL* gene family in pea. (A) The genomic location of the 21 *SPL* genes in pea. (B) Sequence conservation of the SBP domain for all 21 PsSPL. The overall height of each stack indicates the sequence conservation at that position (measured in bits), whereas the height of symbols within the stack reflects the relative frequency of the corresponding amino acid. The two conserved Zn-finger (Zn-1 and Zn-2) structures and the nuclear localisation signal (NLS) are indicated. (http://weblogo.berkeley.edu/logo.cgi).

**Table 1.** List of *Arabidopsis, Medicago* and pea *SPL* genes. The *Medicago truncatula* and pea SPLs were located through BLASTp searches and named according to sequence alignments with *Arabidopsis*. Clade numbering matches the Preston and Hileman paper (2013). Previous numbering of Medicago has been adjusted according to this alignment (see Supplementary Table 2)

| Clade | Arabidopsis |  | Medicago |  | Pea |  |  |  |
| --- | --- | --- | --- | --- | --- | --- | --- | --- |
|  | Name | Reference | Name | Reference | Name | Transcript Reference | Genome Reference | miRNA156 binding site |
| I | AtSPL7 | AT5G18830 | MtSPL7 | Medtr2g020620 | PsSPL7 | PsCam057199 | Psat1g186440 | No |
| II | AtSPL1 | AT2G47070 | MtSPL1a | Medtr1g086250 | PsSPL1a | PsCam048790 | Psat6g160520 | No |
|  | AtSPL12 | AT3G60030 | MtSPL1b | Medtr7g110320 | PsSPL1b | PsCam049132 | Psat3g018160 | No |
|  |  |  | MtSPL1c | Medtr2g046550 | PsSPL1c | PsCam048218 | Psat1g018120 | No |
|  | AtSPL14 | AT1G20980 | MtSPL14 | Medtr1g035010 | PsSPL14 | PsCam048786 | Psat6g132240 | No |
|  | AtSPL16 | AT1G76580 |  |  |  |  |  |  |
| III | AtSPL8 | AT1G02065 | MtSPL8 | Medtr8g005960 | PsSPL8 | PsCam001061 | Psat0s2161g0040 | No |
| IV | AtSPL6 | AT1G69170 | MtSPL6a | Medtr5g046670 | PsSPL6a | PsCam033315 | Psat2g133080 | Yes |
|  |  |  | MtSPL6b | Medtr4g109770 | PsSPL6b | PsCam036737 | Psat4g046440 | Yes |
|  |  |  | MtSPL6c | Medtr2g461920 | PsSPL6c | PsCam025564 | Psat1g127000 | Yes |
| V | AtSPL2 | AT5G43270 | MtSPL2a | Medtr3g085180 | PsSPL2a | PsCam037189 | Psat5g096720 | Yes |
|  | AtSPL10 | AT1G27370 | MtSPL2b | Medtr8g080670 | PsSPL2b | PsCam037577 | Psat7g077080 | Yes |
|  | AtSPL11 | AT1G27360 | MtSPL2c | Medtr8g080680 |  |  |  |  |
|  |  |  | MtSPL2d | Medtr8g080690 |  |  |  |  |
| VI | AtSPL3 | AT2G33810 | MtSPL3a | Medtr2g014200 | PsSPL3a | PsCam033864 | Psat1g203760 | Yes |
|  | AtSPL4 | AT1G53160 | MtSPL3b | Medtr4g088555 | PsSPL3b | PsCam017682 | Psat7g115040 | Yes |
|  | AtSPL5 | AT3G15270 | MtSPL3c | Medtr8g463140 | PsSPL3c | PsCam038909 | Psat4g139840 | Yes |
|  |  |  | MtSPL3d | Medtr2g078770 | PsSPL3d | PsCam047017 | Psat5g242120 | No |
| VII | AtSPL13A | AT5G50570 | MtSPL13a | Medtr3g099080 | PsSPL13a | PsCam046016 | Psat5g052160 | Yes |
|  | AtSPL13B | AT5G50670 | MtSPL13b | Medtr8g096780 | PsSPL13b | PsCam055995 | Psat7g036360 | Yes |
|  |  |  | MtSPL13c | Medtr7g028740 | PsSPL13c | PsCam012994 | Psat3g164360 | Yes |
| VIII | AtSPL9 | AT2G42200 | MtSPL9a | Medtr1g053715 | PsSPL9a | PsCam039166 | Psat6g101960 | Yes |
|  | AtSPL15 | AT3G57920 | MtSPL9b | Medtr7g092930 | PsSPL9b | PsCam037965 | Psat3g069560 | Yes |
| IX | - | - | MtSPL21 | Medtr7g444860 | PsSPL21 | PsCam037459 | Psat3g157680 | Yes |

To assess the conservation of the 21 PsSPL proteins, the 74 amino-acid SBP domain of the pea genes was compared with other published plant SPL sequences (Fig. 2B). All the pea SBP domains contained two zinc-finger motifs (Zn1 and Zn2) and a bipartite nuclear localisation signal (NLS), which acts as a recognition site for binding at the nuclear pore (Robbins et al., 1991; Jans et al., 2000). The N-terminal zinc finger of all the PsSPLs contains three cysteine residues and one histidine (CX_4_CX_16_CX_2_H), except for PsSPL7, which has four cysteine residues (CX_4_CX_16_CX_2_C). The C-terminal zinc finger is conserved across all the PsSPL, with the residue combination of CX_2_CX_3_HX_11_C. These results show that the SBP domain in PsSPL is highly conserved, further validating their function as SPL transcription factors.

### Sequence structure and evolutionary relationships among PsSPLs

To understand the structural diversity of the pea *SPL* family, we examined the exon-intron arrangement for all 21 genes by aligning transcript and genomic sequences and checking intron/exon boundaries against *Medicago SPL* annotations in Phytozome (Goodstein et al., 2012). Fig. 3A shows that the pea *SPL* genes can be classified into two major groups based on length, as either short (<400amino acids) or long (>800 amino acids). The longest genes have ten exons and the shortest only two, but most have 3-4 exons. In all cases, however, the SBP domain is always located at the end of exon 1 across to the beginning of exon 2 (highlighted red in Fig. 3A). Only one significant structural difference was found among the sequences, where the first intron in *PsSPL14* appears to have been lost.

**Figure 3.**
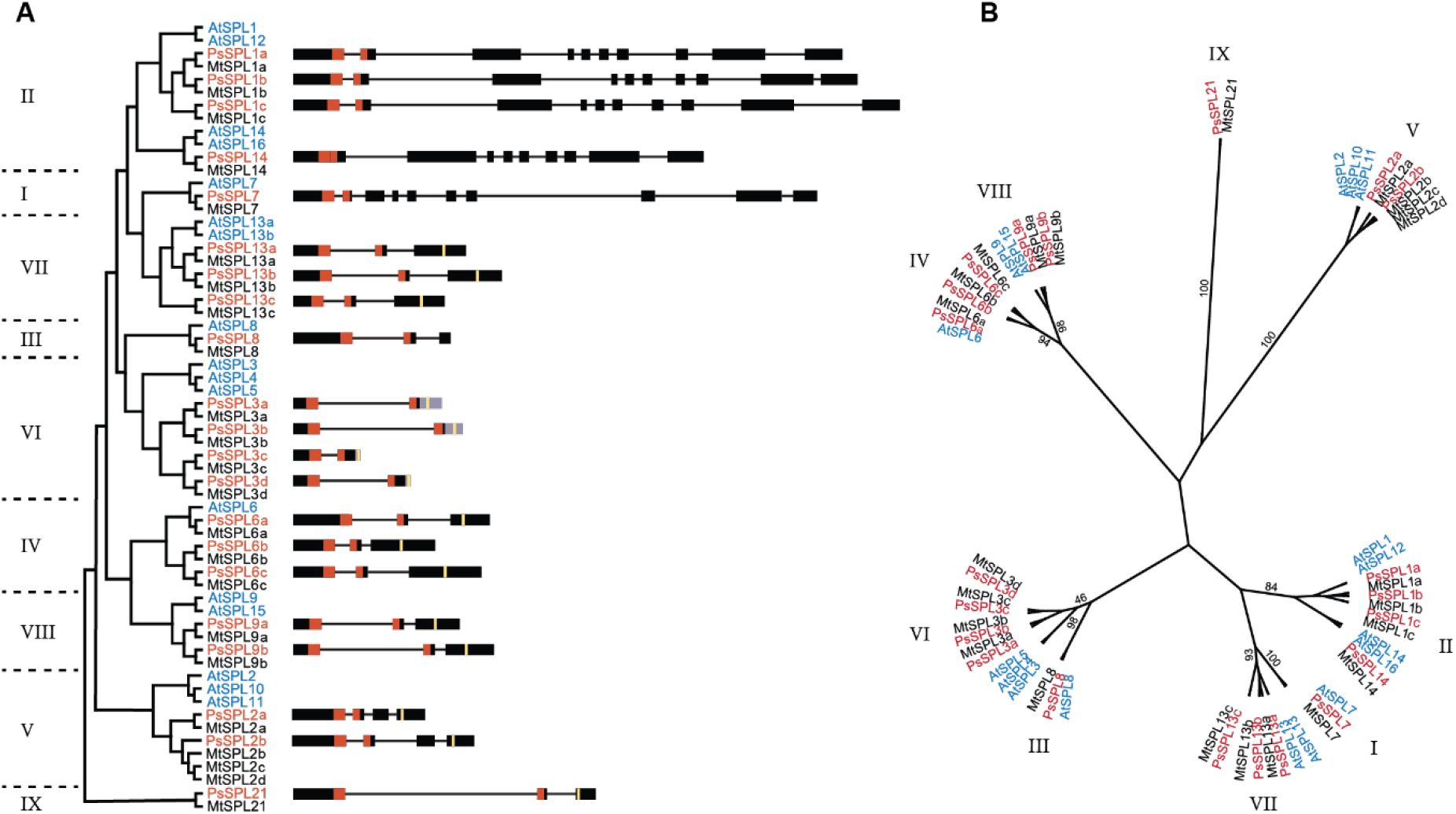
Phylogenetic history of *SPL* genes and their genomic sequence structure in pea. (A) Each of the 21 *SPL* genes in pea has a scaled schematic representation of its exon/intron structure. Structural groupings are represented by a reduced cladogram. The pea and *Medicago* genes are numbered according to their homologues in *Arabidopsis*, except for the legume-specific clade named SPL21. Exons are represented by black boxes, introns by horizontal black lines between the exons. Untranslated 3’ regions (3’UTR) are shaded in grey, the SBP-box domain is shaded red and the *miR156* binding site is represented by a yellow vertical line present within the last exon or 3’UTR. (B) A maximum likelihood tree preformed on the alignment of the SBP-box domain nucleotide sequences from *Arabidopsis thaliana* (At), *Medicago truncatula* (Mt) and *Pisum sativum* (Ps). Bootstrap values obtained from 1,000 trees are indicated for each clade. Clade numbering matches the Preston and Hileman paper (2013). Note, the duplication of SPL13 is present in the Arabidopsis Columbia line used for this study, but most other *Arabidopsis* lines have only a single copy (16 SPL genes in total).

To determine the evolutionary relationships between *Arabidopsis, Medicago* and pea SPL proteins, a maximum likelihood tree was created using the conserved SBP domain nucleotide sequence (222bp). In this analysis, the *Medicago* and pea SPL clustered into nine clades (Numbered I-IX) (Table 1, Fig. 3B). Clades I to VIII grouped with the *Arabidopsis* clades as previously identified by Salinas (2012) and Preston and Hileman (2013), but Clade IX appears to be a distinct clade not represented in *Arabidopsis*. Even though the phylogenetic analysis was performed on the SBP domain only, the groupings also reflect differences in gene structure seen in Fig. 3A. All pea and *Medicago* SPL were named according to the closest relative in *Arabidopsis*. The single sequence for both pea and *Medicago* that did not align with any *Arabidopsis* clade was designated SPL21. Some of the *Arabidopsis* SPL have more than one ortholog in pea and *Medicago* (Table 1, Fig. 3B). These relationships suggest that single ancestral genes duplicated after divergence of Brassicaceae and Leguminosae lineages to generate new paralogs. In pea, 42% of the 21 SPL exist as paralogous pairs and 17% exist as paralogous triplets, indicating significant expansion within the family. A clear example of this is clade IV (SPL6), which is represented in *Arabidopsis* by a single copy, whereas a triplication has occurred in both pea and *Medicago*.

Overall, the number of genes in each clade is relatively variable among the three species, suggesting a complex history of independent duplications and losses in either *Arabidopsis* or the two legumes or in both. For example the AtSPL14/16 duplication is not seen for the legume genes in clade II, but the legume SPL1c, SPL6a/6b and SPL13c genes in clades II, VI and VII are not represented by an *Arabidopsis* ortholog. Clades II, VI, VII and VIII have undergone independent duplications in both species groups, whereas Clade I and III show simple orthology of single-copy genes across the three species (SPL7 and 8). Finally, there was a single instance (MtSPL2c/d) in which additional expansion relative to pea seems to have occurred.

### Expression profiles of SPL genes in pea

The global transcript data generated by Alves-Carvalho *et al*., (2015) has provided spatiotemporal expression profiles for each of the pea *SPL* genes, providing an initial indication of where these genes may be acting during growth and development. We used this publicly available data to create a heat map of expression levels (RPKM) with clustering on the y-axis reflecting similarity of expression pattern (Supplementary Fig. 1). The results show that *SPL* genes are expressed in a wide range of tissues, suggesting diverse functions as seen in other plant species. For example, *SPL6a* is only expressed in the roots and nodules, *SPL3d* is strongly expressed in flowers and *SPL21* is only expressed in the apex. The other genes are grouped on the basis of their preferential expression in the leaves, tendrils, stem and peduncles, or in the apex, flowers and pods, and this grouping mostly aligns with the observed phylogenetic relationships. Clade I (*SPL7*) and II (*SPL1/14*) genes are expressed across all tissue types whereas most of clade III (*SPL8*) and VI (*SPL3*) genes are strongly expressed in apex, flowers and pods. The remaining clades have lower expression levels, but where there is expression, it is also consistently within the apex, flowers and pods. Interestingly, in two cases (*SPL3* and *SPL13*), closely-related paralogs showed marked divergence in their expression.

### The relationship between *miR156/172* and their targets in pea

*SPL* genes that are post-transcriptionally regulated by *miR156* feature a *miR156* binding site motif. To elucidate the potential *miR156*-mediated regulation of *PsSPL* genes, we searched complementary DNA (cDNA) sequences of all *PsSPLs* for the *miR156* mature target site sequence (highlighted in yellow in Fig. 3A; Fig. 4A). We found that all the short *PsSPL* have a *miR156* putative binding site, except for *SPL8*. Genes in Clades III, IV, V, VII, VIII and IX have a *miR156* binding site within the last exon, whereas Clade VI (*PsSPL3*) is unique in that the *miR156* binding site is instead located within the 3’ UTR.

**Figure 4.**
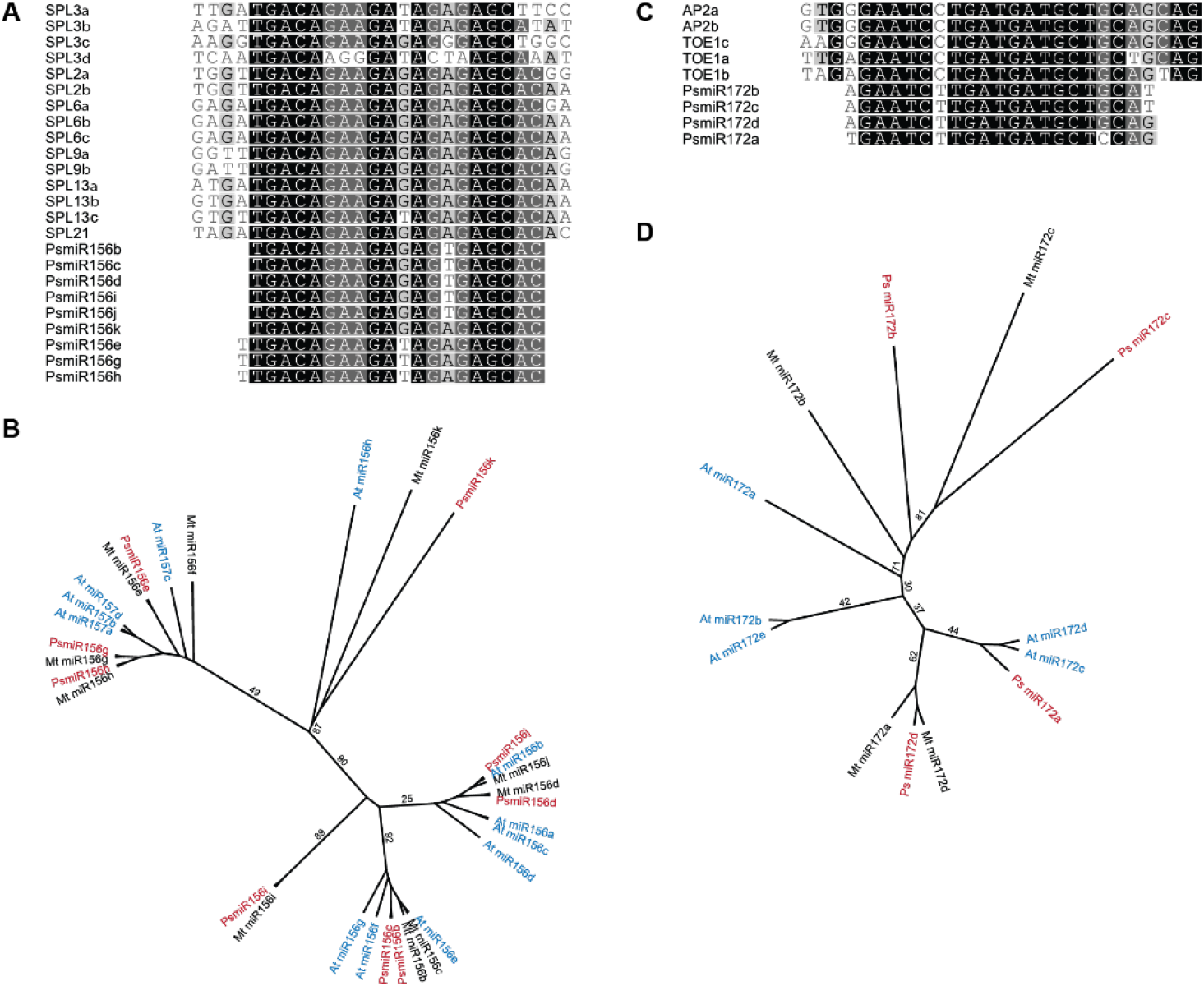
*MiR156* and *miR172* binding sites and evolutionary conservation. (A) Mature *miR156* sequences and the corresponding binding sites in *PsSPL* genes. (B) A maximum likelihood tree preformed on the alignment of the *miR156* precursor sequences from *Arabidopsis thaliana* (At), *Medicago truncatula* (Mt) and *Pisum sativum* (Ps). (C) Mature *miR172* sequences and the corresponding binding sites in *PsAP2-like* genes. (D) A maximum likelihood tree preformed on the alignment of the *miR172* precursor sequences from At, Mt and Ps. Bootstrap values obtained from 1,000 trees.

Plant mature miRNA are known to employ near perfect sequence complementarity to their binding target (Rhoades et al., 2002). This is also the case for the pea *miR156-SPL* association, with the exception of *SPL3a, b, c* and *d*, which have 2, 1, 3 and 7 mismatches respectively in their *miR156* target sites (Fig. 4A). Given the high number of mismatches in *SPL3d*, it is likely that this gene has lost regulation by *miR156* and the fact that it is only expressed in flowers also suggests a specific and divergent function relative to the other *SPL3* genes (Supplementary Fig. 1).

*miR172* targets a subset of the *AP2-like* transcription factors that act to inhibit flowering. We queried the pea genome using the *Arabidopsis* AP2 protein sequence and identified five closely related *AP2-like* transcription factors that all contain the putative *miR172* binding site (Fig. 4C, Supplementary Fig. 2). There are no strict orthologs of AP2 in pea, but two paralogs (PsAP2a/b) are closely related to the AtAP2 and AtTOE3 clade. There is also a pair of pea orthologues corresponding to AtTOE1 (PsTOE1a/b), while the fifth pea gene may be a divergent ortholog of AtTOE2. No sequences clearly orthologous with SMZ and SNZ were identified in either pea or *Medicago*. Further clarity on these relationships will require an expanded analysis including more species and possibly a wider selection of AP2-like family members. However, overall these results confirm that the gene family targets of *miR156* and *miR172* are conserved in pea.

### Analysis of pea *miR156* and *miR172* families

Mature miRNAs are 20-21nt non-coding RNAs that directly affect their target genes by post-transcriptional inhibition or cleavage (Beauclair et al., 2010; Li et al., 2013; Arribas-Hernández et al., 2016). They are first transcribed as a longer RNA sequence (so-called precursor miRNA) that shows self-complementarity, creating a hairpin duplex structure that is then cleaved at both ends to produce the final mature miRNA (Bologna et al., 2012). Sequences of *Medicago miR156* and *miR172* precursor genes were retrieved from miRBase (Kozomara & Griffiths-Jones, 2014; http://www.mirbase.org) and used to identify potential precursor sequences in the pea genome. Nine putative members of the *miR156* family were identified in pea, along with one additional, previously unreported *Medicago miR156* precursor sequence (total 10 in *Medicago*). For *miR172*, four precursor sequences each were identified in *Medicago* and pea, compared to five in *Arabidopsis* (Supplementary Table 3). The pea sequences were named according to the corresponding sequences for *Medicago* given in miRbase.

This analysis also identified several sequences that contained the mature *miR156* sequence but lacked a corresponding duplex sequence. As these would not be able to form a hairpin structure, they were not considered to be valid *miR156* precursors and were excluded. *MtmiR156a* was also excluded, on the basis that the hairpin sequence is unusually long compared with other *Medicago* and *Arabidopsis* precursor sequences and is identical to the sequence of the second intron of *MtSPL21*. It is theoretically possible that the opposing strand of the *SPL21* coding sequence could give rise to the mRNA of a *miR156* precursor sequence, but this is unprecedented and seems unlikely, so we considered *MtmiR156a* to be incorrect. The two *Arabidopsis* precursor sequences *i* and *j* also did not have a matching duplex sequence and were also not included.

Phylogenetic analysis of the *miR156* precursor sequences in *Arabidopsis, Medicago* and pea shows them to group in a manner consistent with the mature *miR156* sequences (Fig. 4A & B). We did not identify a pea ortholog of *MtmiR156f*, suggesting it may have been lost or may still be missing from the genome assembly. Work in *Arabidopsis* has established that *miR156a* and *miR156c* are the primary participants in the age-related pathway and control of vegetative phase change (Yang et al., 2013), and this clade is also represented by two sequences in both *Medicago* and pea; *miR156d* and *miR156j* (Fig. 4B). However, Fig. 4A shows that the same mature *miR156* sequence is also generated by the *PsmiR156b, c* and *i* precursors.

Among the *Arabidopsis miR172* precursors, *miR172a, b* and *d* appear to be critical for flowering, particularly under short-day conditions (Lian et al., 2021; Ó’Maoiléidigh et al., 2021). In this case it was difficult to discern clear relationships between the legume and *Arabidopsis* sequences in the phylogenetic analysis, possibly reflecting the high degree of sequence similarity (Fig. 4D). Nevertheless, *PsmiR172b* appeared to be the pea gene most similar to *AtmiR172a*, and *PsmiR172a* was closest to *AtmiR172d*, but there was no clear pea homolog for *AtmiR172b*.

### Morphological and molecular changes in pea associated with phase change

The pea plant has a highly consistent pattern of ontogenetic change in leaf morphology. The first two alternate nodes produce highly reduced “scale” leaves (Fig. 5A inset), whereas subsequent “true” leaves are pinnately compound and consist of proximal leaflets and distal tendrils (Fig. 5A). The first formed true leaves have a single leaflet pair and simple tendril, but later-formed leaves have increasing numbers of leaflet pairs and paired lateral tendrils, and under long-day conditions, commonly reach a maximum of three pairs at around the first flowering node (Fig. 5B). It has been plausibly suggested that this change in leaf morphology might be a manifestation of VPC (Wiltshire et al. 1994).

**Figure 5.**
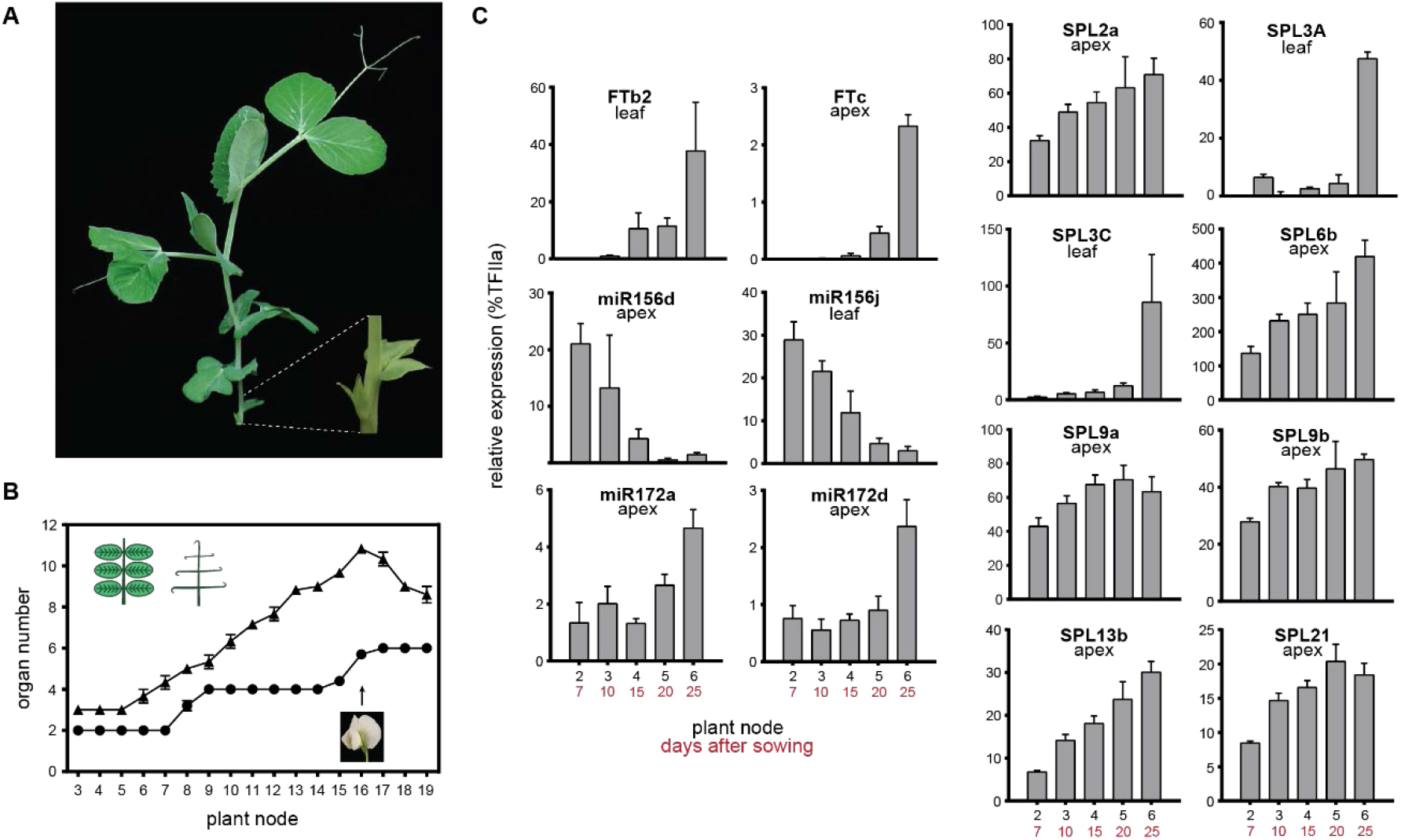
Developmental changes in expression of the *miR156-SPL-miR172* genes in pea. (A) A young pea plant with the scale leaves enlarged for visualisation. (B) The morphological changes in compound leaf development with age in pea grown under long-day conditions. A diagrammatic representation of a leaflet in wild-type pea and tendrils only in an *afila* pea is shown top left. Flowering time is indicated by the arrow and photo. Triangles denote number of tendrils per node (*afila*) and circles denote number of leaflets per node (wild-type). (C) Expression of key developmental genes in wild-type pea (NGB5839). Relative transcript levels were determined in dissected shoot apices or the uppermost fully expanded leaf during development in long days. Values have been normalised to the transcript level of the *TRANSCRIPTION FACTOR II* gene and the error bars represent mean ± the SE for n=3-4 biological replicates, each consisting of pooled material from two plants.

In *Arabidopsis*, VPC is regulated by opposing changes in expression of *miR156, SPL* genes and *miR172*. To examine whether these components display developmental regulation in pea, and whether any in particular show a pattern consistent with a role in VPC, we examined the expression of *miR156* and *miR172* precursor genes, the subset of *SPL* genes with a *miR156* target site, and key *FT* genes (Hecht et al., 2011) (Fig. 5C, Supplementary Fig. 3 & 4). Studies of various mutations within the precursor sequence of *miR156* and *miR172* have shown that the levels of precursor gene expression do quantitatively correlate with mature miRNA expression levels (Werner et al., 2010; Kim et al., 2016; Rojas et al., 2020), and an examination of their expression patterns provides additional detail on their individual contributions.

Among the nine *miR156* precursors, only two (*miR156d* and *miR156j*) showed a clear decrease in expression over the developmental window from 7 to 25 days after sowing, with a 30-fold and 10-fold decrease in expression respectively. These two sequences are not only most similar to the main *miR156* precursor genes in *Arabidopsis* (*AtmiR156a* and *c*) but also were expressed at a much higher levels than all the other pea precursors (with the exception of *miR156c/g* in the leaf; Supplementary Fig. 4). The decrease in their expression could therefore be assumed to contribute to a substantial drop in the level of mature *miR156*. Interestingly, *PsmiR156d* is expressed in shoot apex tissue, whereas *miR156j* is expressed only in the leaf. Among the 21 *PsSPL* genes, six (*SPL2a, 6b, 9a, 9b, 13b* and *21*) showed a pattern of increasing expression in the shoot apex (Fig. 5C). The remaining *SPL* genes that were also expressed in the apex (*2b, 3a, 3c, 6c, 8* and *13a*) did not show an increase in expression with development (Supplementary Fig. 3). Interestingly, two additional *SPL* genes, *SPL3a/3c* showed increasing expression in the mature leaf at the end of the time series. Of the four *miR172* genes only two, *miR172a* and *miR172d*, showed an increase in expression, with the greatest increase between day 20 and day 25. These two genes were expressed only in the apex and at relatively low levels. However, all the *SPL* and *miR172* genes that showed an age-dependent increase were already substantially expressed even in seven-day-old seedlings when the key age-regulated *miR156* genes are most strongly expressed.

In an earlier study, we showed that the induction of the key pea florigen (*FT*) genes in LD occurs between 10 and 15 days after sowing, and coincides with the physiological commitment to flower (Hecht *et al*. 2011). The results in Fig. 5C are consistent with this, and suggest that if there is a distinct transition from a juvenile to an adult vegetative stage, it must occur prior to this point. Expression of *miR156* genes is significantly reduced by two weeks old and the expression of both *SPL* and *miR172* seems to be relatively high at this stage too. Combined, these results suggest that the juvenile phase in this WT pea variety may be very short and any switch to the mature vegetative phase occurs over the first week after emergence.

## Discussion

The so-called “aging” or phase change pathway, involving the sequential *miR156-SPL-miR172-AP2* interaction, is now thought to be broadly conserved in flowering plants (Poethig, 2013; Wang and Wang, 2015). However, it has not yet been explored extensively in many major plant groups, including legumes, and has not been examined at all in pea, despite its long-standing status as a genetic and physiological model and an important cool-season legume crop. Here we have defined all four gene families in pea and examined certain key aspects of their regulation in relation to developmental changes in leaf morphology, with the aim of assessing the potential for these genes to contribute to an aging response, and how such a response might be manifested.

Our genome-level analyses confirm that clear that pea and related legumes possess a full array of aging pathway genes. The recently released draft pea genome (Kreplak et al., 2019) includes 21 SPL genes representing all of the eight previously-described clades and one additional divergent gene (*PsSPL21*) with no apparent *Arabidopsis* ortholog (Fig. 3). Of the 21 pea *SPL* genes, 14 contain a functional *miR156* binding site, with the majority located within the third exon, except for the *SPL3* clade where an imperfectly complementary site is located in the 3’ UTR. In *Arabidopsis*, similar 3’ UTR binding sites have been functionally validated using transgenic *SPL* plants either sensitive or resistant to *miR156* (Xu et al., 2016), suggesting that *PsSPL3a-c* should also be regulated by *miR156*.

Six of the pea *SPL* genes show a pattern of significant increase in expression in the shoot apex during development, broadly consistent with observations of *SPL* genes implicated in VPC in other species (Xie, 2006; Salinas et al., 2012; Xu et al., 2016), while two others (*PsSPL3a* and *PsSPL3b*) with increasing transcript levels were expressed predominantly in leaves. In *Arabidopsis* six of the ten *SPL* genes with a *miR156/7* target sequence act with partial redundancy in regulation of VPC and floral induction (Xu et al., 2016). Interestingly, apart from the divergent *PsSPL21*, the other pea genes that show increasing expression are all co-orthologs of these *Arabidopsis* genes. In several cases the pattern of increasing expression is not seen shared between paralogs (e.g. *PsSPL2a* vs. *PsSPL2b*), suggesting that the presence of the *miR156* target site alone is not enough to specify increased expression, and that genomic context may be more important than phylogeny for whatever actually does determine it.

Among *miR156* precursors, only two (*PsmiR156d* and *PsmiR156j*) showed a characteristic decline in expression during development and these, along with *PsmiR156g*, were also the most strongly expressed, by at least an order of magnitude relative to all others. In *Arabidopsis*, both of the main precursor genes *AtmiR156a* and *AtmiR156c* are expressed mostly at the shoot apex (Fouracre and Poethig, 2019) and the same was true for *PsmiR156d*. Interestingly, *PsmiR156j* was strongly expressed in mature leaf tissue, with comparatively low levels in apex tissue and it is possible that *miR156j* works in conjunction with *SPL3a/c* in leaf related processes. It is also plausible that it may contribute to a pool of mature *miR156* acting at the shoot apex, considering that the mature *miR156* sequence is small enough to be transported long distances through the plant and there is evidence to suggest that this can happen (Martin et al., 2009; Kasai et al., 2010; Marín-González and Suárez-López, 2012; Bhogale et al., 2014). Overall, these results suggest that in pea, the key interactions related to VPC are likely to involve *miR156d/j* and *SPL2a/3a/3c/6b/9a/9b/13b/21*.

Considering *SPL* targets, we identified four *miR172* precursor genes and a clade of five *AP2-like* homologues that all contain the conserved *miR172* binding site. Interestingly, in pea, although Ps*miR172d/j* did show a significant increase in expression by day 25, this occurred later than for *SPL* genes and after the induction of *FT* genes in the leaf and apex, suggesting that it may have only occurred in tissues already specified as reproductive. Five of the six Arabidopsis *AP2-like* genes are important targets of *miR172* and contribute substantially to regulation of flowering time through regulation of *FT* and of other genes acting upstream and downstream of *FT* (Mathieu et al., 2009; Ó’Maoiléidigh et al., 2021). Whereas pea and *Medicago* have two clear *TOE1* orthologs, they have only one other *TOE* gene (*PsTOE1c*) of uncertain relationship to *AtTOE2, SNZ* and *SMZ*. In addition, recent evidence on the function of this third *TOE1* gene in pea and its ortholog in *Lotus japonicus* indicates a function largely restricted to regulation of floral organ number and symmetry (Weng et al., 2020). This suggests that unlike in *Arabidopsis*, the pea *miR172* genes may not be directly regulated by SPL proteins, and also that miR172 and its targets may not act substantially to repress the floral transition, which has already occurred by around 14 days under these conditions (Fig. 5C; Hecht *et al*., 2011). However, one alternative interpretation is that *miR172* regulation may be highly tissue-specific and any differences may be obscured in analysis of larger, more complex tissue samples. It may also be that under long days in pea other pathways influencing flowering are more dominant than any effect of *miR172* system, given that the effect of *miR172* on flowering in *Arabidopsis* is greater under short day than long days (Lian et al., 2021).

The nature of VPC in pea is not clear. The bipinnate pea leaf shows a characteristic increase in leaflet number as the plant ages and previous studies (e.g. Wiltshire *et al*. 1994) have attempted to frame this in terms of phase change. However, although leaflet number by definition increases in a step-wise pattern with the addition of leaflet pairs, and it is tempting to consider each increase as a distinct phase, it is likely that this pattern may instead just reflect some threshold for leaflet initiation against a background of a more general increase in the growth and/or determinacy of successive leaf primordia. This more gradual tendency is illustrated in the *afila* mutant (Goldenberg, 1965) in which leaflets are essentially replaced by tendrils (Fig. 5B), and suggests that the juvenile-adult transition might not be associated with a specific morphological marker.

The expression patterns we observed for the *miR156* and *SPL* genes also have implications for understanding the nature of VPC in pea. The developmental changes in expression of the key *miR156* and *SPL* genes between 7 and 15 days and the fact that all six developmentally-regulated *SPL* genes in the apex are already substantially expressed 7 days after sowing suggests that VPC is already underway at this point and an even earlier time point might be necessary to fully define any critical window, if one indeed exists. Interestingly, apart from the increase in leaflet numbers, the only other marked change in pea leaf morphology occurs early in development, with the transition from vestigial scale-like leaves at the first two nodes to a true foliage leaf at node 3. It is not yet clear whether this transition might also be a manifestation of VPC, but it is shared with other related temperate legumes that have epigeous germination, it does not vary across the pea germplasm, and there is little, if any genetic evidence indicating that it can be substantially altered. Significantly, the developmental fate of leaves at the first few nodes is already established in the mature pea seed, suggesting that if the scale-leaf to true-leaf transition is indeed a manifestation of VPC, then causal molecular changes would actually occur while the seed is still developing. It may therefore be interesting in future to examine expression of *miR156* and *SPL* genes in the embryonic shoot during seed development and immediately following germination.

It is clear that a definitive analysis of VPC pathway roles in pea will require impairment or manipulation of its *miR156* and *SPL* components. However, some evidence for the influence of this pathway is slowly accumulating across several different legume species. Overexpression of *miR156* precursors in soybean, *Medicago* and *Lotus* have some common features, including delayed flowering and increased branching, biomass and rate of development. In soybean, overexpression of *GmmiR156b* also increased seed and pod size, but had no effect on plant height (Cao et al., 2015; Sun et al., 2019). In contrast to soybean, overexpression did influence plant height in *Medicago sativa* (alfalfa) (Aung *et al*., 2015a/b; Gao *et al*., 2016) and actually reduced leaf, seed and pod size in *Medicago truncatula* (barrel clover) (Wang et al., 2019). In *Lotus japonicus*, transgenic plants with ectopic expression of *LjmiR156a* showed underdeveloped roots and reduced nodulation along with delayed flowering and increased branching (Wang *et al*., 2015). In soybean, loss-of-function mutants have been generated for all four *SPL9* genes (Bao et al., 2019), and in both *Medicago sativa* and *Medicago truncatula*, for *SPL8* (Gou et al., 2018). In all cases, the phenotypic changes were similar to those reported for the *miR156* overexpression plants, with additional effects on salt and drought tolerance documented in alfalfa. Evidence in other non-legume species also suggests that the *miR156/SPL* system can influence leaf physiological function even in the absence of major morphological changes (Lawrence *et al*., 2021b). It may therefore be that in pea (and other species lacking clear heteroblasty) aging is primarily reflected in physiological differences and does not directly relate to either the scale leaf or the leaflet number transitions.

In this study, our overview of *miR156-SPL* pathway components and their expression in pea has identified those that are most likely to have a significant role and provides the necessary and important groundwork for future study of these genes aimed at clarifying their roles in growth and development. Results from functional studies in other legume species suggest that these roles are wide-ranging and impact morphological and physiological features critical for crop performance and yield. These results also imply that in pea, certain genes in the pathway may be detectable in a forward genetic approach targeting a range of phenotypes such as flowering time, branching and developmental rate. At least one such pea mutant (*aero1*) has been described, in which the leaflet number change is accelerated (Taylor and Murfet, 2003), but the molecular identity of this gene is so far unknown. A complementary forward genetic approach targeting altered timing of the leaflet number transition would also be potentially useful to help clarify the mechanistic basis of this trait and the extent to which it might be related to *miR156/SPL* function.

## Materials and Methods

### Identification of *SPL* genes in pea

We performed a BLAST search (Altschul et al., 1990) against the pea transcriptome and genome with the known Arabidopsis and Medicago *SPL* genes found previously in NCBI, as queries to determine pea *SPL* sequences and their location in the pea genome. To exclude the redundant sequence of *PsSPL* genes, we aligned all sequences with CLUASTALW (Chenna et al., 2003) in Geneious 8.1.9 (https://www.geneious.com). All protein sequences of putative *PsSPL* genes were manually checked for the SBP domain and aligned using WebLogo 3 (Crooks et al., 2004). The intron-exon structures of *PsSPL* genes were determined by mapping cDNAs to genomic sequence and were illustrated proportionally to the lengths of introns and exons.

### Identification of *miR156* and *miR172* genes in pea

Known *miRNA156* and *miR172* sequences in Medicago and Arabidopsis from MIRBase (Kozomara and Griffiths-Jones, 2014) were BLASTed against the pea genome to determine the pea homologous sequences. All miRNA precursors were validated by determining if they had the mature miRNA sequence and duplex sequence through alignment and stem-loop structure using miRNAFold (Tav et al., 2016). According to the previously characterised miRNAs in plants (Rhoades et al., 2002; Mallory and Vaucheret, 2006), genes that contained sequences with fewer than three mismatches to the mature *miR156* sequence were considered to be putative targets of *PsmiR156* and *PsmiR172* in pea.

### Phylogenetic analysis

For phylogenetic analyses, the genes in Arabidopsis, Medicago and pea were aligned using ClustalW and a maximum-likelihood tree using the HKY85 substitution model was performed on the alignments using PHYML v2.2.3 (Guindon et al., 2010) in Geneious v8.1.9. Bootstrap values were obtained from 1000 trees. The trees were edited for clarity using Adobe Illustrator (Adobe Inc. 2020; https://adobe.com/products/illustrator).

### Expression profile and heatmap

Expression levels (RPKM) for each pea SPL was collated from the publicly available pea transcriptome data published by Alves-Carvalho *et al*. (2015). This data was used to analyse the expression profiles of PsSPL genes in 12 tissue types (root, nodule, shoot, leaf, tendril, stem, peduncle, apical node, flower, pods, embryo, seed). If there were multiple values for a tissue type, they were pooled and the mean value used. Hierarchical clustering of the expression profiles of SPL genes was performed using R (R Core Team, 2017).

### Plant growth and conditions

The wild-type pea (Pisum sativum) line NGB5839 used for this research is a dwarf derivative of ‘Torsdag’ carrying the le-3 and hr mutations (Lester et al., 1999; Weller et al., 2012). All plants were grown in a 1:1 mixture of dolerite chips and vermiculite topped with potting mix and received nutrient solution weekly. Plants were grown in growth cabinets at 20°C under 150 μmol m^-2^ s^-1^ white light from cool-white fluorescent tubes (18hr light/6hr dark).

### Gene expression studies

Harvested tissue used in RT-qPCR experiments (Fig. 4, Supp Fig. 3) consisted of both leaflets from a fully expanded leaf or dissected apical buds (~2 mm) from two plants. Samples were frozen in liquid nitrogen and total RNA extracted using the SV Total RNA Isolation System (Promega). RNA concentrations were determined using a NanoDrop 8000 (Thermo Scientific). Reverse transcription was performed in 20mL with 1mg of total RNA using the Tetro cDNA synthesis kit (Bioline). RT-negative (no enzyme) controls were included to monitor for genomic DNA contamination. RT-qPCR reactions using SYBR green chemistry (SensiFast; Bioline) were set up with a CAS-1200N robotic liquid handling system (Corbett Research) and run for 50 cycles on a Rotor-Gene Q (Qiagen). Two technical replicates and three biological replicates were performed for each sample. Values have been normalised to the transcript level of the *TRANSCRIPTION FACTOR II* gene. Primer details are included in Supplemental Table 4.

## Acknowledgements

Many thanks to the glasshouse staff, Tracey Winterbottom and Michelle Lang for looking after the plants. Thanks to Jakob Butler for creating the heatmap in R. We acknowledge the use of facilities administered by the UTAS Central Science Laboratory. This work was supported by the Australian Research Council (J.L.W.)

## Author Contributions

J.L.W., J.K.V.S. and V.H. conceived and designed the experiments. J.K.V.S. performed the experiments. G.A. and J.B contributed early release sequence data. J.K.V.S., J.L.W. and V.H. analysed the data. J.K.V.S and J.L.W. wrote the article.

