## Supplementary Figures & Tables for "Defining the components of the miRNA156-SPL-miR172 aging pathway in pea and their expression relative to changes in leaf morphology"

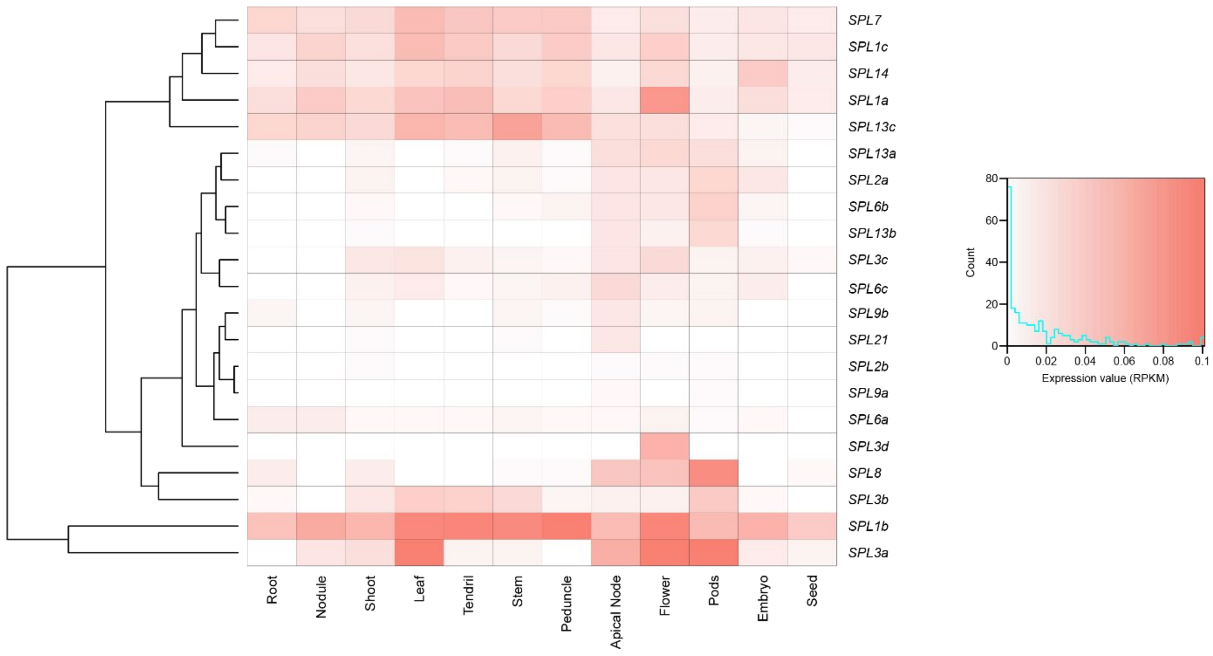


**Supplementary Figure 1. Tissue-specific expression heat map for pea *SPL* genes**

Expression levels for each of the pea *SPL* genes within different tissue types. Data was taken from the publicly available pea transcriptome data previously published by Alves-Carvalho *et al*. (2015). Average expression levels (RPKM) were calculated for root, nodule, shoot, leaf and seed. White = no expression, Red = highest expression levels. Clustering on the y-axis reflects similarity of expression pattern.

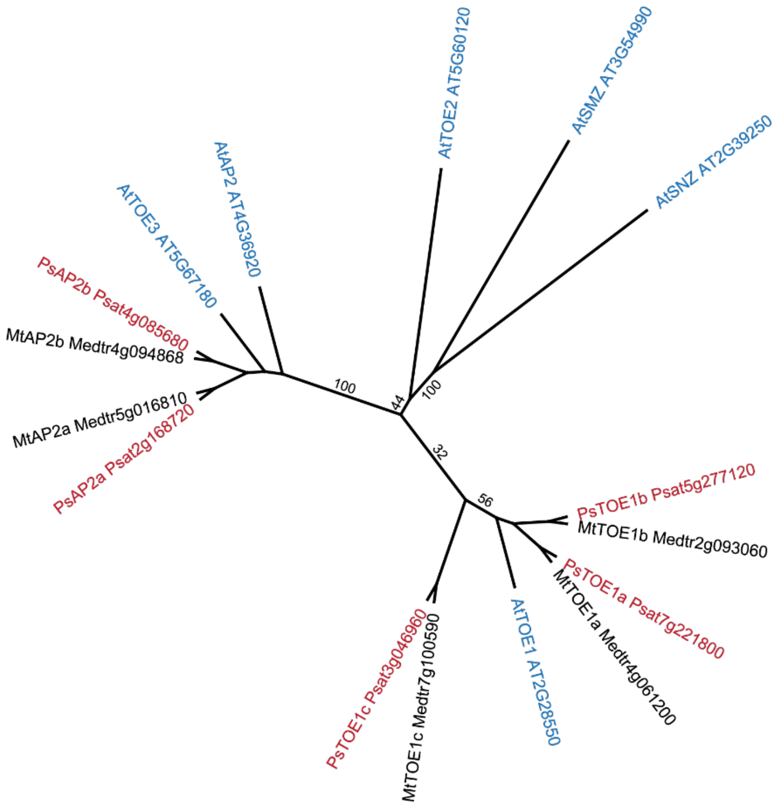


**Supplementary Figure 2.** ***AP2-like*** **gene family**

*Arabidopsis*, *Medicago* and pea AP2-like protein sequences were aligned and phylogenetic analysis performed using maximum-likelihood software within Geneious (PhyML). Bootstrap values were obtained from 1000 trees.

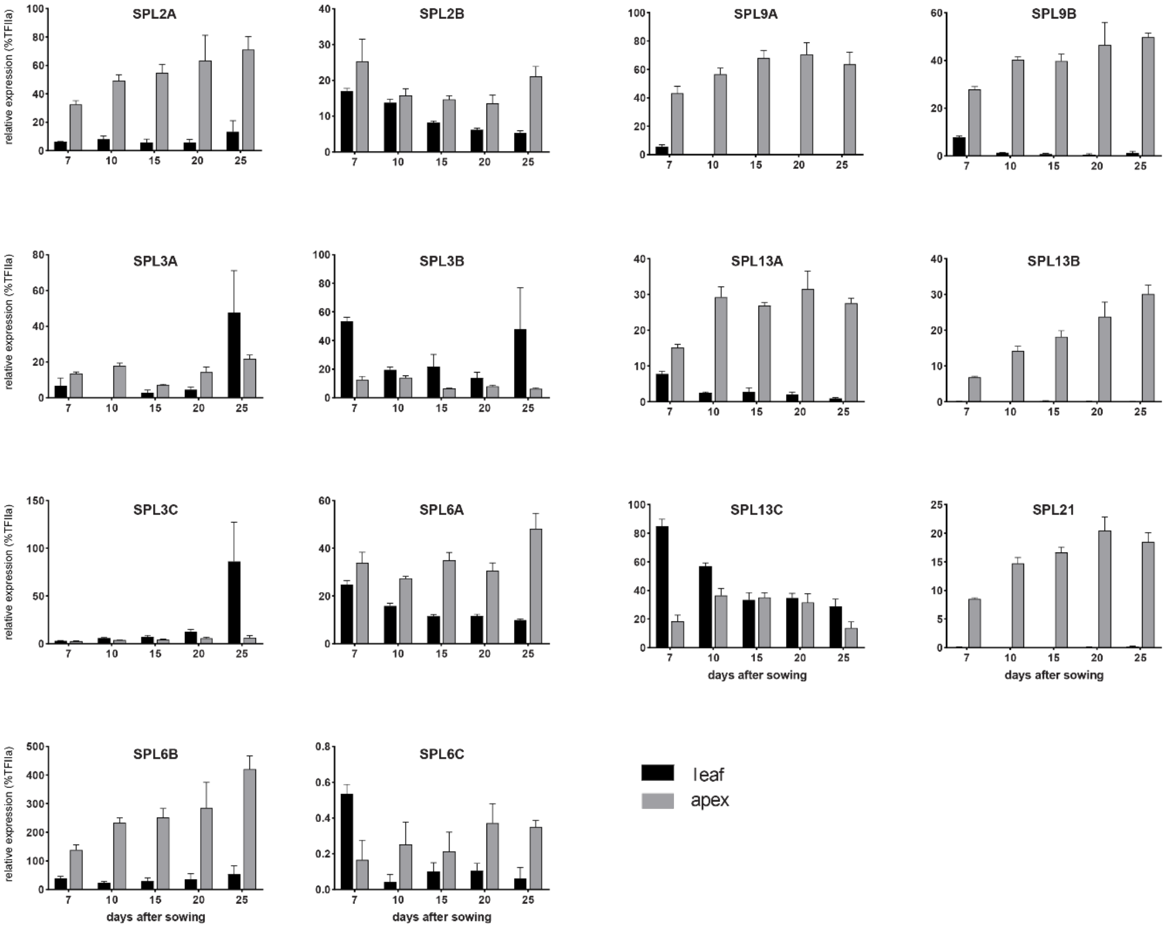


**Supplementary Figure 3. Expression of *SPL* genes in pea**

Expression of all *miR156*-regulated *SPL* genes in a developmental series in wild-type pea (NGB5839). Relative transcript levels were determined in dissected shoot apices or the uppermost fully expanded leaf during development in long days. Values have been normalised to the transcript level of the *TRANSCRIPTION FACTOR II* gene and the error bars represent mean ± the SE for n=3-4 biological replicates, each consisting of pooled material from two plants.

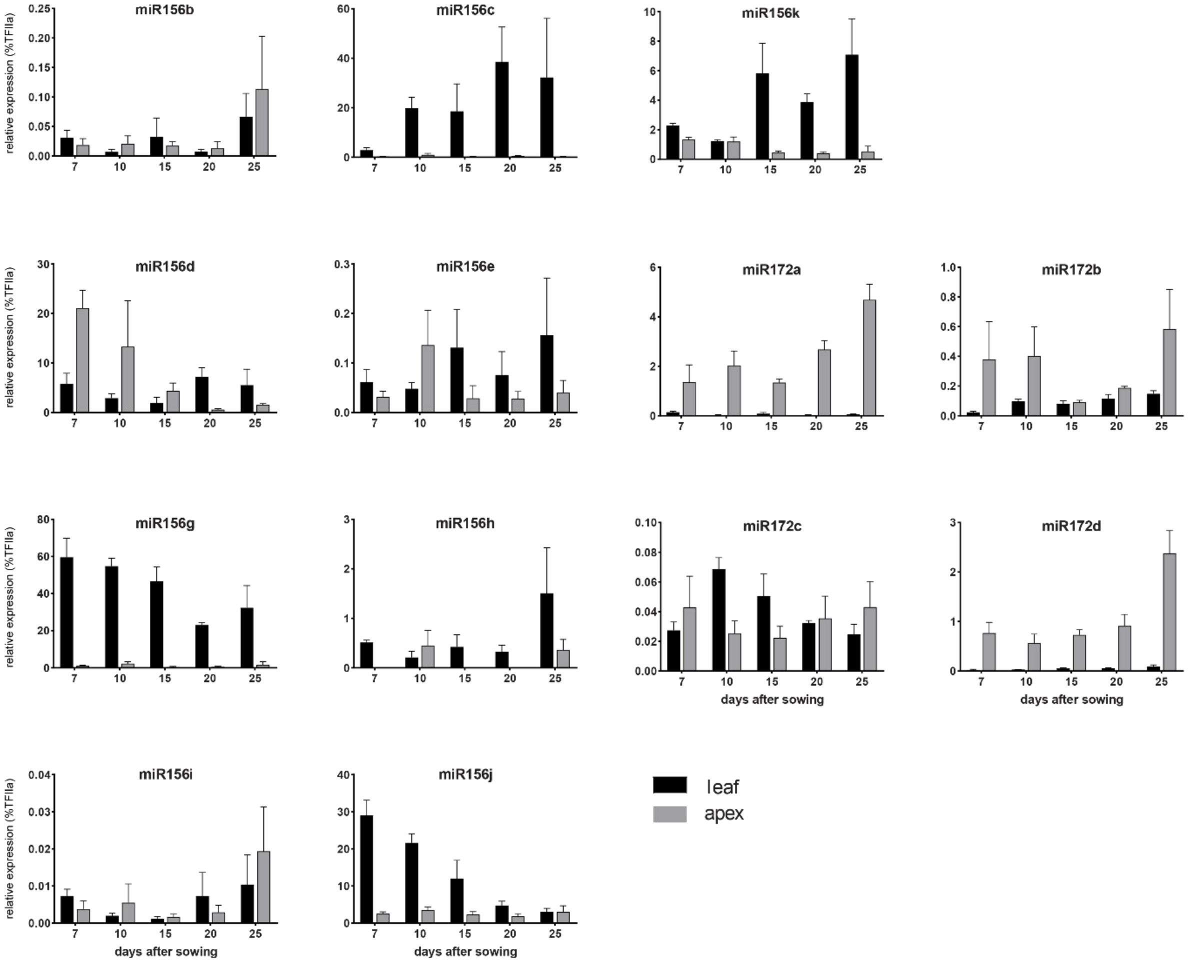


**Supplementary Figure 4. Expression *miRNA156* and *miR172* precursors in pea**

Expression of *miR156* and *miR172* genes in a developmental series in wild-type pea (NGB5839). Relative transcript levels were determined in dissected shoot apices or the uppermost fully expanded leaf during development in long days. Values have been normalised to the transcript level of the *TRANSCRIPTION FACTOR II* gene and the error bars represent mean ± the SE for n=3-4 biological replicates, each consisting of pooled material from two plants.

### Supplementary Table 1. List of *Medicago truncatula SPL* genes

List of names given to the complete set of *Medicago truncatula SPL* genes and their previous names from other papers. Accession and genomic numbers are given. Paper author and year of publication are at the top of each column (Preston & Hileman, 2013; Aung *et al* . 2015b; Gao *et al* ., 2016; Wang *et al* ., 2019).

| Name (this paper) | Accession | Number | Preston & Hileman 2013 | Aung 2015 | Gao 2016 | Wang 2019 |
|---|---|---|---|---|---|---|
| | | | maximum likelihood | none | unknown | neighbour-joining |
| | | | SBP-domain only | | SBP-domain only | full length CDS |
| SPL1a | XP_003626036 | Medtr7g110320 | SPL1a | | SPL1a | SPL16 |
| SPL1b | XP_003591325 | Medtr1g086250 | SPL1b | | SPL1b | SPL1 |
| SPL1c | XP_013463701 | Medtr2g046550 | | | | SPL12 |
| SPL2a | XP_003601767 | Medtr3g085180 | SPL2 | SPL12 | SPL2 | SPL10b |
| SPL2b | XP_013446576 | Medtr8g080670 | | | | SPL11a |
| SPL2c | XP_013446577 | Medtr8g080680 | | | | SPL11b |
| SPL2d | XP_013446578 | Medtr8g080690 | | | | SPL10a |
| SPL3a | XP_003593617 | Medtr2g014200 | | | | SPL3a |
| SPL3b | XP_013456994 | Medtr4g088555 | | | | SPL3b |
| SPL3c | XP_01344549 | Medtr8g463140 | | | | SPL5 |
| SPL3d | XP_013464662 | Medtr2g078770 | | | | SPL4 |
| SPL6a | XP_003614226 | Medtr5g046670 | SPL6 | SPL6 | SPL6 | SPL6b |
| SPL6b | XP_013458014 | Medtr4g109770 | | | | SPL6c |
| SPL6c | XP_013464103 | Medtr2g461920 | | | | SPL6a |
| SPL7 | XP_003594035 | Medtr2g020620 | | | | SPL7 |
| SPL8 | XP_003626693 | Medtr8g005960 | SPL8 | | SPL8 | SPL8 |
| SPL9a | XP_013467649 | Medtr1g053715 | | | | SPL9 |
| SPL9b | XP_003625236 | Medtr7g092930 | SPL9 | | SPL9 | SPL15 |
| SPL13a | XP_003602795 | Medtr3g099080 | SPL14a | SPL13 | SPL14 | SPL13b |
| SPL13b | XP_013446991 | Medtr8g096780 | | | | SPL13a |
| SPL13c | XP_013447951 | Medtr7g028740 | | | | SPL13c |
| SPL14 | XP_003589683 | Medtr1g035010 | | | | SPL14 |
| SPL20 | XP_013448322 | Medtr7g444860 | | | | SPL2 |

**Supplementary Table 2. Synteny between pea (Ch4) and Medicago (Ch8) in region of *SPL8***

*Medicago* genes in the region of *SPL8* were sequentially BLASTed against the pea genome to find their location and infer the position of *PsSPL8* .

| Gene | Medicago | Pea | Pea Location |
|---|---|---|---|
| PA domain | Medtr8g005800 | Psat4g209320 | Chromosome 4 |
| remorin | Medtr8g005810 | Psat4g209360 | |
| CCR4 | Medtr8g005820 | Psat4g209400 | |
| PPR | Medtr8g005870 | Psat4g208920 | |
| zinc-finger | Medtr8g005880 | Psat4g208900 | |
| matrixin | Medtr8g005900 | Psat4g209040 | |
| SPL8 | Medtr8g005960 | Psat0s2161g0040 | Scaffold 02161 |
| CMD | Medtr8g005980 | Psat4g208760 | Chromosome 4 |
| isomerase | Medtr8g005990 | Psat4g208800 | |
| transporter | Medtr8g006050 | Psat4g227000 | |
| transporter | Medtr8g006090 | Psat4g227040 | |
| hydrolase | Medtr8g006370 | Psat4g226960 | |
| ERF | Medtr8g006410 | Psat4g226920 | |
| DnaJ | Medtr8g006430 | Psat4g226880 | |

**Supplementary Table 3. List of *Arabidopsis* , *Medicago* and pea *miR156* and *miR72* sequences**

List of miR156 and miR172 sequences found in miRbase (http://www.mirbase.org) and the matching pea sequences located in the genome. Some have been excluded from phylogenetic and qPCR analysis for reasons identified in the comment column. Differing bases within the mature sequence are highlighted in red. Accession details are given where available.

| Arabidopsis | | | |
|---|---|---|---|
| **Name** | **Accession** | **mature** | **Comment** |
| AtMIR156a | MI0000178 | TGACAGAAGAGAGTGAGCAC | |
| AtMIR156b | MI0000179 | TGACAGAAGAGAGTGAGCAC | |
| AtMIR156c | MI0000180 | TGACAGAAGAGAGTGAGCAC | |
| AtMIR156d | MI0000181 | TGACAGAAGAGAGTGAGCAC | |
| AtMIR156e | MI0000182 | TGACAGAAGAGAGTGAGCAC | |
| AtMIR156f | MI0000183 | TGACAGAAGAGAGTGAGCAC | |
| AtMIR156g | MI0001082 | CGACAGAAGAGAGTGAGCAC | |
| AtMIR156h | MI0001083 | TGACAGAAGAAAGAGAGCAC | |
| AtMIR156i | MI0019232 | TGACAGAAGAGAGAGAGCAG | no duplex, excluded |
| AtMIR156j | MI0019234 | TGACAGAAGAGAGAGAGCAC | no duplex, excluded |
| AtMIR157a | MI0000184 | TTGACAGAAGATAGAGAGCAC | |
| AtMIR157b | MI0000185 | TTGACAGAAGATAGAGAGCAC | |
| AtMIR157c | MI0000186 | TTGACAGAAGATAGAGAGCAC | |
| AtMIR157d | MI0000187 | TGACAGAAGATAGAGAGCAC | |
| **Medicago** | | | |
| **Name** | **Accession** | **mature** | **Comment** |
| Mtr-MIR156a | MI0001752 | TGACAGAAGAGAGTGAGCAC | SPL21 match, excluded |
| Mtr-MIR156b | MI0005571 | TGACAGAAGAGAGTGAGCAC | |
| Mtr-MIR156c | MI0005584 | TGACAGAAGAGAGTGAGCAC | |
| Mtr-MIR156d | MI0005585 | TGACAGAAGAGAGTGAGCAC | |
| Mtr-MIR156e | MI0005595 | TTGACAGAAGATAGAGAGCAC | |
| Mtr-MIR156f | MI0005605 | TTGACAGAAGATAGAGAGCAC | |
| Mtr-MIR156g | MI0005607 | TTGACAGAAGATAGAGGGCAC | |
| Mtr-MIR156h | MI0005613 | TTGACAGAAGATAGAGAGCAC | |
| Mtr-MIR156i | MI0005626 | TGACAGAAGAGAGTGAGCAC | |
| Mtr-MIR156j | MI0018375 | TGACAGAAGAGGGTGAGCAC | |
| Mtr-MIR156k | chr3:23939450..23939545 | TGACAGAAGAGAGAGAGCAC | this study |
| **Pea** | | | |
| **Name** | **Location** | **mature** | **Comment** |
| Psa-MIR156a | chr3:304502722..304502795 | TGACAGAAGAGAGTGAGCAC | SPL21 match, excluded |
| Psa-MIR156b | chr6:16607671..16607765 | TGACAGAAGAGAGTGAGCAC | |
| Psa-MIR156c | scaffold02479:91960..92056 | TGACAGAAGAGAGTGAGCAC | |
| Psa-MIR156d | chr5:61818177..61818268 | TTGACAGAAGAGAGTGAGCAC | |
| Psa-MIR156e | ch7:9428774..9428664 | TTGACAGAAGATAGAGAGCAC | |
| Psa-MIR156f | - | no match | |
| Psa-MIR156g | ch1:51411459..51411591 | TTGACAGAAGATAGAGAGCAC | |
| Psa-MIR156h | ch3:194758065..194757972 | TTGACAGAAGATAGAGAGCAC | |
| Psa-MIR156i | super-scaffold6646:352044..352136 | TGACAGAAGAGAGTGAGCAC | |
| Psa-MIR156j | ch6:19401893 to 19401984 | TGACAGAAGAGAGTGAGCAC | |
| Psa-MIR156k | ch2:150678564 to 150678470 | TGACAGAAGAGAGAGAGCAC | |

| Arabidopsis | | | |
|---|---|---|---|
| **Name** | **Accession** | **mature** | **Comment** |
| AtMIR172a | MI0000215 | AGAATCTTGATGATGCTGCAT | |
| AtMIR172b | MI0000216 | AGAATCTTGATGATGCTGCAT | |
| AtMIR172c | MI0000991 | AGAATCTTGATGATGCTGCAG | |
| AtMIR172d | MI0000992 | AGAATCTTGATGATGCTGCAG | |
| AtMIR172e | MI0001089 | GGAATCTTGATGATGCTGCAT | |
| **Medicago** | | | |
| **Name** | **Accession** | **mature** | **Comment** |
| Mtr-MIR172a | MI0005600 | AGAATCCTGATGATGCTGCAG | |
| Mtr-MIR172b | MI0018368 | AGAATCTTGATGATGCTGCAT | |
| Mtr-MIR72c | MI0025218 | AGAATCTTGATGATGCTGCAT | |
| Mtr-MIR72d | MI0018369 | AGAATCTTGATGATGCTGCAG | |
| **Pea** | | | |
| **Name** | **Location** | **mature** | **Comment** |
| Psa-MIR172a | ch2:67576896..67576977 | TGAATCTTGATGATGCTCCAG | |
| Psa-MIR172b | ch5:543143995..543143896 | AGAATCTTGATGATGCTGCAT | |
| Psa-MIR172c | ch7:258006569..258006437 | AGAATCTTGATGATGCTGCAT | |
| Psa-MIR172d | sacffold02449:253180..253279 | AGAATCTTGATGATGCTGCAG | |

**Supplementary Table 4. List of primers used in qRT-PCR experiments**

| Gene | ID | Primer | Sequence | Temp | Lenth (bp) |
|---|---|---|---|---|---|
| SPL2A | Psat5g096720 | SPL2A-1F | CATCATCTCTTTCTTCATCGCCC | 60 | 64 |
| | | SPL2A-1R | CTCGGAAAAGCAAATGGGCC | | |
| SPL2B | Psat7g077080 | SPL2B-1F | ATGGAAGTGGTTGCTCTGGG | 60 | 190 |
| | | SPL2B-1R | ACAGAAGAAAGCTCCGGTGC | | |
| SPL3A | Psat1g203760 | SPL3A-1F | TGAGGGAAAGAGAAGCTATGAGT | 60 | 185 |
| | | SPL3A-1R | GCTTACCACCACTTAGATCAGC | | |
| SPL3B | Psat7g115040 | SPL3B-1F | TTTGAAGAGGAGGAGGAAGG | 61 | 212 |
| | | SPL3B-1R | ATCTCTGCAATGTGTACGGC | | |
| SPL3C | Psat4g139840 | SPL3C-1F | CTGTTGTGATGGTTGCAGGG | 61 | 191 |
| | | SPL3C-1R | AGGTTGATGTTGTCCTTTGCC | | |
| SPL6A | Psat2g133080 | SPL6A-1F | CGCACTTCCACTTTACCTTCC | 61 | 209 |
| | | SPL6A-1R | CGCTTATCATCATCGAACTCAGC | | |
| SPL6B | Psat4g046440 | SPL6B-1F | GTGAATTTGATGATGGTAAGCGC | 61 | 161 |
| | | SPL6B-1R | AGATGCTTGTGGTGGCTTGG | | |
| SPL6C | Psat1g127000 | SPL6C-1F | ATTCACTCAGGAAGGGCTGG | 60 | 185 |
| | | SPL6C-1R | CTAGATCCGGTTCCGTCTTCG | | |
| SPL9A | Psat6g101960 | SPL9A-1F | GAGCTCCGTGTTTACCTCGC | 60 | 155 |
| | | SPL9A-1R | TTGTTTGATCGCCGCGTACC | | |
| SPL9B | Psat3g069560 | SPL9B-1F | CCAACTCTCTCTTAACTTCGCG | 60 | 156 |
| | | SPL9B-1R | CGGGTGTTGTGGTTTGATTTCC | | |
| SPL13A | Psat5g052160 | SPL13A-1F | ACTTCATTCCAAGACTCCTGAGG | 60 | 162 |
| | | SPL13A-1R | TCAGGCTGAGGTTTTCTTCTCC | | |
| SPL13B | Psat7g036360 | SPL13B-1F | GCAGGAAACGTTTAGATGGGC | 60 | 175 |
| | | SPL13B-1R | ACATCAGTTCCACCCCAAGTAG | | |
| SPL13C | Psat3g164360 | SPL13C-1F | TCCTGTTGTGTTGGTTGGAGG | 60 | 153 |
| | | SPL13C-1R | AGAGAAGGTGGTTGAGGTTTCC | | |
| SPL21 | Psat3g157680 | SPL20-1F | TAGGAGAAGACTTGCAGGGC | 60 | 156 |
| | | SPL20-1R | GCACTGCACCTTACGGAGAG | | |
| FTb2 | Psat3g206880 | FTLe-2-F7 | CGACTACCGGGACAGCATTT | 62 | 186 |
| | | FTLe-2-R7 | CAGGTGAACCAAGGTTATAAAC | | |
| FTc | Psat3g091040 | FTL-L1-8F | GATATTCCAGCCACAACAAGC | 62 | 149 |
| | | FTL-L1-7R | TTATGACGCCACTCTGGAGCAA | | |
| TFIIa | Psat5g118600 | TFIIa-qF | GCAACCTCCTTCTCCTTGGAT | 60 | 75 |
| | | TFIIa-qR | TCTTCCCGTCCTTCCACATAA | | |

| Gene | Primer | Sequence | Temp | Lenth (bp) |
|---|---|---|---|---|
| miR156b | PsMIR156B-1F | GCATGTCTGGTTCAATCGAGAG | 60 | 163 |
| | PsMIR156B-2R | TGGCAGAAAACAGAACAGTAAATAG | | |
| miR156c | PsMIR156C-1F | TTAAGGGTAAGGGCGGTGAC | 61 | 122 |
| | PsMIR156C-1R | AGAGAGATGGTGGTGGGGTG | | |
| miR156d | PsMIR156D-3F | CTCCTTTCATATCCGAGAATCTTTC | 65 | 189 |
| | PsMIR156D-3R | TGAATGGGAGCATAACAGGAC | | |
| miR156e | PsMIR156E-3F | CTGAGGAGTGGAGGAATTAATAGG | 60 | 171 |
| | PsMIR156E-3R | GTAGAGGAAAATGGGAATTGG | | |
| miR156g | PsMIR156G-1F | TGTTTAAGAGGGAGAGGGAGGA | 60 | 175 |
| | PsMIR156G-1R | GAGCACAAAGGAAATGAAATGCA | | |
| miR156h | PsMIR156H-1F | TTGCCATGAGAGGTTGAGGC | 60 | 137 |
| | PsMIR156H-1R | TGCATGAGAAGGGTGATGGTG | | |
| miR156i | PsMIR156I-1F | AATCCACAGAGAGAGAGAGAGACATAA | 60 | 155 |
| | PsMIR156I-1R | GAGAGACACATTCGAGAGATGA | | |
| miR156j | PsMIR156J-1F | CCATATCTAGTGTGTTTTCTGTTG | 65 | 183 |
| | PsMIR156J-1R | GAGAAATTTCCTTTCCAGCAC | | |
| miR156k | PsMIR156K-1F | TGGCAAGAAGGGTTGAATTGG | 60 | 185 |
| | PsMIR156K-1R | GGCAAAGGTGGAAAGTCTAGTG | | |
| miR172a | PsMIR172A-1F | TCTGGCGGCGTTGATTAG | 60 | 137 |
| | PsMIR172A-1R | CGAGATCTGATGAAAACAGTCG | | |
| miR172b | PsMIR172B-1F | TGATGATCATAGTCGTTGTTTGC | 60 | 145 |
| | PsMIR172B-1R | CTTGGATTTATAAAGTCGTTTATGG | | |
| miR172c | PsMIR172C-1F | CGTCATTTGCGGATGTAGC | 60 | 165 |
| | PsMIR172C-1R | AAACTATTTTTAAAGCCGTCTATGG | | |
| miR172d | PsMIR172D-1F | AACAGTCAATGTTTGCTAGTGG | 60 | 137 |
| | PsMIR172D-1R | TGAAGCTGTATTAGTCATTGATTGC | | |
